# Emotional Distractors Capture Attention even at Very Low Contrast Levels: ERP evidence

**DOI:** 10.1101/2024.06.05.597626

**Authors:** Germán A. Cipriani, Dominique Kessel, Fátima Álvarez, Uxía Fernández-Folgueiras, Manuel Tapia, Luis Carretié

## Abstract

Emotional visual stimuli, whether appealing or aversive, preferentially capture exogenous attention due to their evolutionary significance. This study assessed whether such capacity persists at low contrast levels, where stimuli are minimally perceived. To this end, we recorded behavioral and electrophysiological (event-related potentials, ERPs) indices of attentional capture from 38 participants who were exposed to negative, neutral, and positive scenes, each presented at four distinct contrast levels. These contrast levels had previously resulted in a correct recognition rate of up to 25%, 50%, 75%, and 100% in a previous sample of 235 participants. Participants were presented with these scenes as distractors while simultaneously performing a perceptual task involving line orientation discrimination. The ERP results confirmed the expected emotional effect on exogenous attention and, critically, unveiled its persistence across all contrast levels. Specifically, occipito-parietal P1 (88-119 ms) was larger for negative than for positive distractors, while in a spreaded N2 component, positive distractors elicited larger amplitudes relative to both negative (213-354 ms) and neutral (213-525 ms) images. These findings reinforce the advantage of emotional distractors in accessing neural processing automatically and highlight the existence of a temporal negativity bias. Importantly, our novel findings emphasize the robustness of this exogenous attention pattern, present even under limited perceptual conditions.

## 1. INTRODUCTION

Evolutionary pressure has made emotional stimuli, both appetitive and aversive, especially efficient at automatically capturing attention (for a review see Carretié, 2014). At the behavioral level, this attentional capture, also termed exogenous attention, is reflected in slower reaction times and decreased accuracy when emotional stimuli are presented as distractors in an ongoing task than when neutral distractors are presented (e.g., Carretié et al., 2013b; Schönwald & Müller, 2014; Sussman et al., 2013). At the neural level, indexes of exogenous attention may also be found in event-related potentials (ERPs). Thus, enhanced amplitudes to emotional distractors (as compared to neutral), may be observed in several ERP components, particularly in P1 (e.g., Carretié & Ruiz-Padial, 2016; Carretié et al., 2004, 2005, 2009, 2019; Fernández-Folgueiras et al., 2022; Soares et al., 2017), the anterior P2 (e.g., Carretié et al., 2004, 2011; Feng et al., 2012; Trauer et al., 2012), and the family of N2 components -i.e., N2x- (e.g., Buodo et al., 2010; Carretié et al., 2004, 2019; Eimer & Kiss, 2007; López-Martín, 2013).

Previous studies (e.g., Carretié et al., 2004; 2005; 2008; 2011; 2013b; Hinojosa et al., 2009; López-Martín et al., 2013; Mercado et al., 2006; 2009) were mostly focused on the modulation of exogenous attention by high-level variables and properties of distractors (e.g., their category —words, faces or scenes— and emotional valence and arousal), as well as by factors related to the ongoing task (e.g., level of difficulty, cognitive nature) or to the individual (e.g., anxiety level). However, more recently, there has been growing interest in understanding how low-level parameters affect exogenous attention to emotional stimuli. Some of the analyzed parameters have been the spatial location of the distractors, their spatial frequencies, their durations, their subliminal condition, their chromaticity, the figure-ground iso/hetero luminance, or the environmental luminosity (e.g., Carretié et al., 2012, 2013a, 2016; 2017, 2022; Soares et al., 2017). Critically, it has been shown that several factors associated with magnocellular visual processing, such as visual periphery, figure-ground heteroluminance, low spatial frequencies, or motion, benefit attentional capture by emotional stimuli (e.g., Alorda et al., 2007; Carretié et al., 2007, 2012, 2017; Miskovic et al., 2015; Pourtois et al., 2005). In this respect, an important factor that may modulate exogenous attention to emotional stimuli is the contrast level.

In real-life scenarios, contrast is strongly variable as a function of general luminosity and the presence -or not- of shadows propitiated by a strong light source (usually the sun). Given that the magnocellular visual processing system is more sensitive to (even low) contrast levels than the parvocellular (Merigan & Maunsell, 1993; Purpura et al., 1988; Tootell et al., 1988, Tootell & Nasr, 2017), it is reasonable to hypothesize that emotional stimuli should be able to induce an attentional capture even at very low contrast levels. To our knowledge, the only study exploring contrast effects on attention to emotional stimuli was developed by Lee and colleagues (2010), who presented fearful and neutral facial expressions as targets under two contrast levels, “high” and “low”. Participants were asked to endogenously attend faces in order to assess the expression in each trial. The classification task was correctly performed (i.e., more hits than misses) in both high and low contrast levels, although accuracy diminished in the latter. At the ERP level, fearful expressions elicited larger EPN and LPP amplitudes than neutral, both typically indexing endogenous attention, regardless of the contrast level. Therefore, the preferential endogenous attention to emotional stimuli seems to survive at low contrast levels. However, several open issues remain. First, given that exogenous attention is controlled by -at least partially-different neural mechanisms (e.g., Corbetta et al., 2008) and serves different functional scopes (e.g., Carretié, 2014) than endogenous, it is necessary to test the effect of contrast level in the former type of attention. Second, and taking into account that even the low-contrast level in the aforementioned study was apparently easy to recognize, the question arises on whether clearly poor contrast conditions are still compatible with enhanced attention to emotional stimuli. Third, whether the contrast effects on attention to negative stimuli, usually more critical in adaptive terms (e.g., Ekman, 1992; Öhman et al., 2000), are generalizable to positive/ appetitive stimuli is a relevant but unexplored issue. Fourth and finally, in comparison to faces, scenes are effective in inducing extreme emotional reactions, and they can elicit larger neurophysiological responses (e.g. N2x or LPP) (e.g. Mavratzakis et al., 2016; Thom et al., 2014; but see Carretié et al., 2013b). In this regard, from an evolutionary perspective, it is fundamental to understand how non-facial emotional distractors (such as scenes) capture attention in low contrast conditions.

Based on this rationale, the aim of the present study was to assess the influence of contrast levels on exogenous attention to emotional scenes which presented positive, negative and neutral valence. In order to explore this issue, we employed a concurrent but distinct target-distractor (CDTD) task (Carretié, 2014), which consists of the simultaneous presentation of physically segregated targets and distractors. Participants are instructed to endogenously attend the first, while ignoring the latter. In our case, emotional scenes were the distractors, while two bars, whose orientation participants had to discriminate, were presented as targets. As indicated, behavioral and/ or neural indices show greater interference when emotional distractors (either positive or negative) compared to neutral ones, are presented: at the behavioral level, larger error rates and slower reaction times are observed, while, at the neural level, larger P1, P2, and/or N2x amplitudes are evidenced. Here, distractors were presented with four contrast levels, which were correctly identified at different rates (%) by an independent sample: i) a control condition with no contrast change (0, 76-100% accuracy); ii) a medium contrast level (229, 51-75%); iii) a low contrast level (247, 25-50%); and iv) a very low contrast condition (250, <25%). Both behavioral (reaction times and error rates) and neural indices (ERPs) of attentional capture were measured during the CDTD task. No data-driven hypotheses may be formulated given the absence of previous literature involving the variables we are exploring (i.e., exogenous attention, negative and positive emotional valence, emotional scenes, an extremely low contrast condition).

## 2. METHODS

### 2.1. Participants

Forty-four volunteers participated in this experiment, although data from only 38 of them could eventually be analyzed to guarantee a minimum number of trials per condition, as explained later (32 women, age range of 18–36 years, mean=20.34, standard deviation [SD]=3.17). The study had been approved by the Ethics Committee of the Universidad Autónoma de Madrid. All participants were students of Psychology, provided their informed consent according to the Declaration of Helsinki, and received academic compensation for their participation. They reported normal or corrected-to-normal visual acuity.

### 2.2. Stimuli and procedure

Participants were placed in an electrically shielded, sound-attenuated, and video-monitored room, at a distance of approximately 85 cm from the screen. Participants performed a CDTC task (Figure 1). Stimuli were presented on a VIEWpixx® (120 Hz) screen using Psychtoolbox task programming extensions for Matlab (Brainard, 1997; Kleiner, Brainard & Pelli, 2007). Each trial was composed of two bars that served as targets of the main task, placed over a background distractor picture. Thus, targets consisted of two blue bars presented on both sides of a blue fixation dot and showing variable orientations. The orientations of both bars were either the same or different (50%-50%; in the latter case, whatever their orientations, the difference was always 18°). Bar orientations appeared in random order but, in sum, the same combinations of bars were presented in all conditions. The visual angle of the whole stimulus was 41.1° (width) × 23.8° (height), and the size of each bar was 0.84º (width) × 2.7º (height). Background stimuli employed as distractors consisted of 60 emotional images selected from *EmoMadrid: emotional picture database for affect research* (Carretié, Tapia, López-Martín & Albert, 2019). The selection of images (20 Negative, 20 Neutral, and 20 Positive) was made using a MatLab® Script. In this way, selected stimuli were significantly different in both Valence [*F*(2,57) = 506.6, *p*<0.001] and Arousal [*F*(2,57) = 103.1, *p*<0.001] dimensions. Post-hoc contrasts indicated that Negative and Positive pictures showed different Valence [*p* < 0.001] but not different Arousal levels (*p* > 0.05), and that they differed from Neutral pictures in both dimensions [all *p* < 0.001]. Furthermore, selected pictures were balanced in spatial frequency, luminosity, and physical complexity [all *p*>0.05]. Then, all images were digitally altered to be presented as gray-scaled and at four different levels of contrast using Adobe® Photoshop® CS3 Extended.

**Figure 1.**
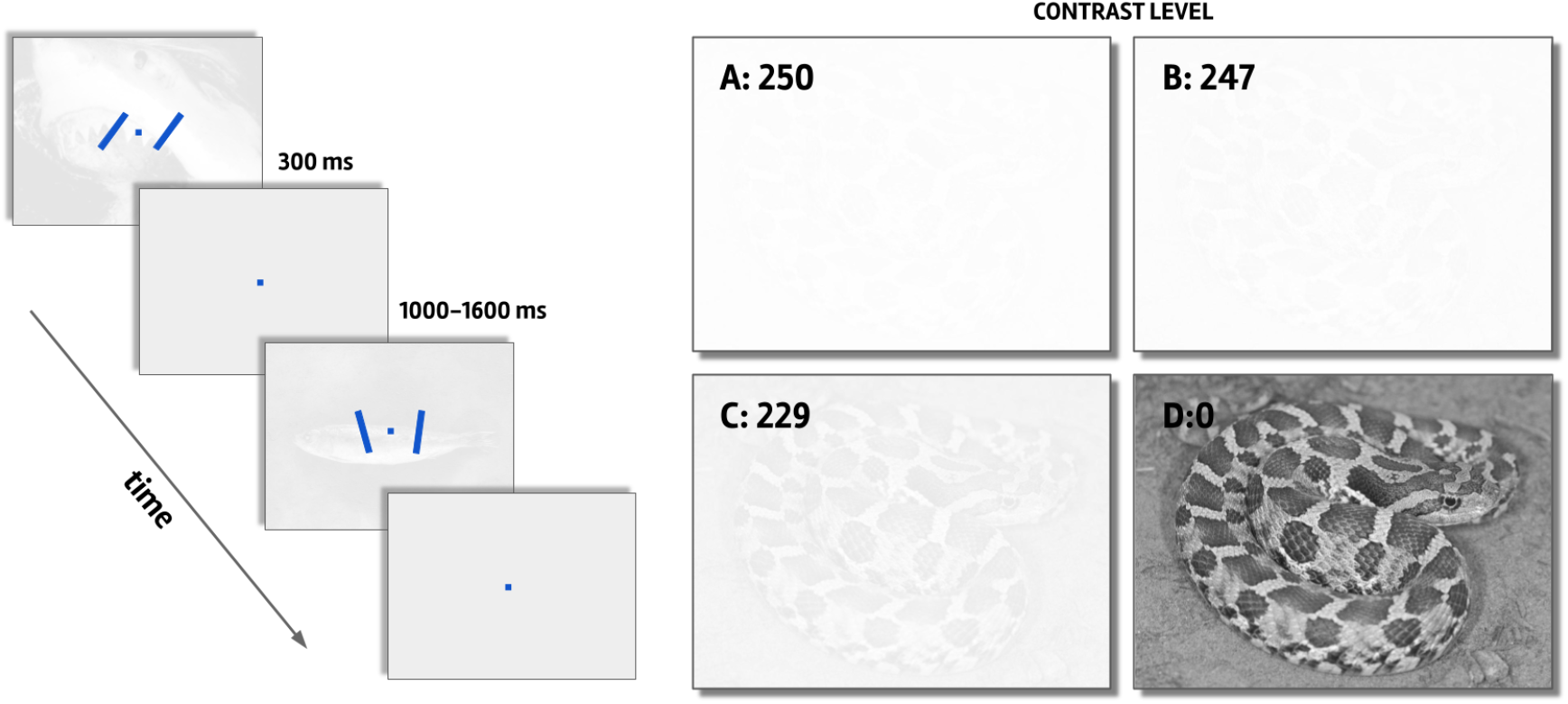
Experimental task. On the left, the concurrent but distinct target-distractor task is shown. Participants were asked to report whether both blue bars were equally oriented (first example) or not (second example), while emotional background scenes were manipulated as negative (distractor in the first example), neutral (distractor in the second example) or positive at different contrast levels (229 in the example series). On the right, the four contrast levels for emotional images are shown: A, 250 (very low); B, 247 (low); C, 229 (medium); and D, 0 (unaltered or intact).

In order to determine and select the four perceptually critical contrast levels of the images, an independent sample of 235 participants (204 women, age range of 18–31 years, M=19.24, SD=1.76) evaluated 32 images at eight contrast levels generated using the function ‘Levels’ in Photoshop, which modifies the files with brightness values ranging from 0 (0%=black) to 255 (100%=white). Participants were asked to identify the content of each image through a computerized questionnaire by choosing one of two different descriptions presented along with the image that better described it. Correct descriptions were created as an agreement between several researchers, while the incorrect one was randomly chosen from the descriptions of the rest of the images. The data collected from this sample allowed to select the four levels of contrast alteration: i) 250 or very low (less than 25% of the sample identified the images); ii) 247 or low (between 25% and 50%); iii) 229 or medium (between 51% and 75%); 0 or normal (between 76% and 100%). Contrasts levels were created as follows: in the 0 contrast level, the image remained unchanged; in the 229, 247, and 250 contrast levels, luminance values were proportionally redistributed within a range that started from 229 (89.8%), 247 (96.9%), and 250 (98%), respectively, and that ended in 255 (100%). For each level, the average rooted-mean square contrast -RMS-values (Kukkonen et al., 1993) across images were: 0.2557 (SD=0.0656) for 0; 0.0261 (SD=0.0067) for 229; 0.0078 (SD=0.0023) for 247; and 0.0052 (SD=0.0013) for 250. Pairs of paired t-test comparisons were calculated (Holm-Bonferroni adjusted), and all RMS values significantly differed between contrasts [all p < .001].

All stimuli were displayed on the screen for 300 ms, followed by a blue fixation dot on a blank gray screen of a random duration between 1000 and 1600 ms, being the resulting mean stimulus onset asynchrony 1600 ms. Participants were asked to look continuously at the center of the screen, to press ―as accurately and rapidly as possible― one key if both bars presented the same orientation, and a different key if they did not. Before starting the experiment, they completed a practice block of ten trials (five of each bar orientation) to ensure they understood the task instructions. Thus, there were 40 trials of each condition (250, 247, 229, 0 × Negative, Neutral, Positive), consisting of 20 different pictures, each of them presented two times within the four different contrast levels, which were presented in blocks. The total number of 480 trials was displayed in four blocks of 120, separated by a rest period. In all cases, the experiment started presenting the block of pictures with the lowest contrast level and ended with the highest contrast level, in order to avoid a facilitating effect during the lower contrast conditions due to a previous presentation of the same picture showing a higher contrast (i.e., being better recognizable).

### 2.3. Recording and preprocessing

Continuous electroencephalographic (EEG) activity was recorded using the ActiveTwo BioSemi system (BioSemi, Amsterdam, The Netherlands). Sixty-four electrodes were placed on the scalp following the distribution of the International 10-20 System. Electrooculographic (EOG) data were recorded supraorbitally and infraorbitally (vertical EOG), as well as from the left versus right orbital rim (horizontal EOG). The EEG signal was pre-amplified at the electrode. The sampling rate was 512 Hz, and a low-pass fifth order CIC filter with a −3 dB cutoff point at 104 Hz was employed. Following the BioSemi design, the voltage at each active electrode was recorded with respect to a common mode sense (CMS) active electrode and a Driven Right Leg (DRL) passive electrode, replacing the ground electrode. Behavioral data were recorded through a common keyboard.

Offline analyses were conducted using Fieldtrip software (http://fieldtrip.fcdonders.nl; Oostenveld et al., 2011). Electrophysiological data were re-referenced offline to the nose tip, and an offline digital Butterworth bandpass filter (fourth order zero phase two-pass forward and reverse) of 0.3 - 30 Hz was applied using Fieldtrip software. The continuous recording was divided into 1000 ms epochs for each trial, beginning 200 ms before stimulus onset. Baseline correction was performed based on this 200 ms pre-stimulus recording. The inevitable lag between the marks signaling stimulus onsets (or “triggers”) in EEG recordings and its actual onset on the screen was measured employing a photoelectric sensor as described in https://www.youtube.com/watch?v=0BPwcciq8u8, and corrected during preprocessing.

Ocular artifact removal was conducted through an Independent Component Analysis based strategy (Jung et al., 2000), as implemented in Fieldtrip. After this process, a second stage of visual inspection of EEG data was conducted, in order to remove any possible remaining interference. Outlier trials (responses outside the range of mean ± 3 standard deviations, and responses before 200 ms), incorrect trials, and trials with no response were also eliminated. This combination of automatic and manual rejection procedures led to the average number of 29.6, 29.6, 29.2, 30.8, 31.1, 31.2, 30.7, 30.6, 30.9, 29.0, 29.6, and 30.2 trials (respectively for 250, 247, 229, 0 × Negative, Neutral, Positive; SD: 4.6, 5.1, 4.9, 4.1, 4.0, 3.8, 3.6, 3.7, 4.1, 4.5, 4.7, and 4.8). The minimum number of trials accepted for averaging was 20 trials per participant and condition. Noisy electrodes were recovered through interpolation from neighbor channels up to 10% of the total number of electrodes.

### 2.4. Data analysis

#### 2.4.1. Behavioral data

All analyses were performed using R (version 3.6.1, Team, 2013). Outlier trials were defined as described above. Generalized linear mixed models (GLMM) were applied on reaction times (RT) and on the binary accuracy response (correct or incorrect) for each task, employing the *glmer* function (lme4 package; Bates et al., 2009). GLMMs are recommended for RT (Lo & Andrews, 2015) and the analysis of dichotomous variables (Dixon, 2008) because there are no assumptions of variance homogeneity and normal distribution of residuals. Additionally, the use of GLMMs avoids variable transformations, which may lead to misinterpretations of results (Whelan, 2008). To establish the best structure for the random and fixed components we followed a well-known procedure (see Zuur et al., 2009). First, several random structures were proposed for each GLMM, from those we selected the one that yielded the lowest Akaike information criteria (AIC). We compared random structures including all possible combinations of the random slopes of the fixed effects and their interactions at the participant level. Subsequently, the fixed structure was selected by comparing pairs of models with likelihood ratio tests (see Supplementary Table 1); the first one was the full model which included Contrast (250, 247, 229, 0), Valence (Negative, Neutral, Positive), and their interactions. Afterwards, we systematically dropped -one by one-non-explicative interactions or effects based on the one that yielded the lowest deviance. For the accuracy GLMM, the best random structure included Contrast as a random slope within participants, while for the GLMM assessing RT, a Valence slope was also included. For the RT model, an inverse gaussian with an identity link function was used, while for the binary accuracy response a binomial distribution with a logit link function was selected. Finally, for each model, an analysis of deviance (analogous to ANOVA) was performed. For significant main and interaction effects, post-hoc z-tests adjusted by the Holm-Bonferroni method were calculated on least-square means.

#### 2.4.2. ERP data

To assess potential attentional capture effects, ERP data was analyzed through a Mass Univariate Analyses (MUA) approach (Groppe et al., 2011) using the *FclustGND* function from the latest extension of the MUA, the Factorial Mass Univariate Toolbox (FMUT; Fields, 2017). This data-driven approach allows the assessment of ERP components at the whole scalp, while controlling for the type I error, and largely avoiding the visual selection of time windows and electrodes (Fields & Kuperberg, 2020). This method consists in performing an ANOVA at each time point and electrode of interest; then, clusters are defined as adjacent time points at the temporal level and electrodes at the topographic level, showing effects surpassing a given threshold (p < 0.01); finally, all F values are summed to compose the cluster mass statistic (Summed F). Permutations are executed to estimate the null distribution of the cluster mass, which is used to calculate a p-value for each cluster. Considering the previous research on exogenous attention effects to emotional stimuli on ERP components (Carretié, 2014) we were interested in P1, P2, and N2 components. On the ERP averages at each electrode the three components were observed (Supplementary Figure 1), though the temporal latency of the P2 component largely varied depending on the topographical region (fronto-central peak: 170-215 ms; occipito-parietal peak: 210-250 ms) and it partially overlapped with the N2 component. Therefore, two separate windows were selected to perform MUAs: the first latency (50-150 ms) assessed P1, while the second one assessed P2 and N2 (150-600 ms). ERP data were downsampled from 512 to 128 Hz. In case of a significant Valence or Contrast x Valence interaction cluster, post-hoc Bonferroni-corrected MUA paired two-tailed t-test contrasts on difference waves were calculated for the significant electrodes componing the largest time window in the cluster (*clustGND* function). For the purposes of the present study, we were not interested in Contrast clusters, so they will be reported but not subjected to further analysis.

## 3. RESULTS

Individual behavioral and ERP data, as well as supplemental information mentioned above, are openly available at https://osf.io/e54q7/. In the case of ERPs, data are provided in the form of a four-dimension matrix with a size of 64 EEG recording channels × 513 data points x 12 conditions × 38 participants.

### 3.1. Behavior

Analyses on RTs yielded a global average of 636 ms; least-square means for each condition are shown in **Table 1**. The best GLMM included *Contrast* as a fixed effect, but not *Valence*. The *Contrast* main effect was significant [Wald χ2(3)=278.3, p<.001]. Corrected post-hoc z-tests revealed that RTs were slowest in the 0 (M=666ms, SE=5ms) and 250 (M=660ms, SE=8ms) contrasts [0 vs 229, z=13.2, p<.001; 0 vs 247, z=10.6, p<.001; 229 vs 250, z=8.4, p<.001; 247 vs 250, z=9.7, p<.001; 0 vs 250, z=0.9, p=.39], and fastest in the 229 (M=607ms, SE=6ms) and 247 (M=611ms, SE=7ms) contrasts [229 vs 247, z=.8, p=.407].

**Table 1.**
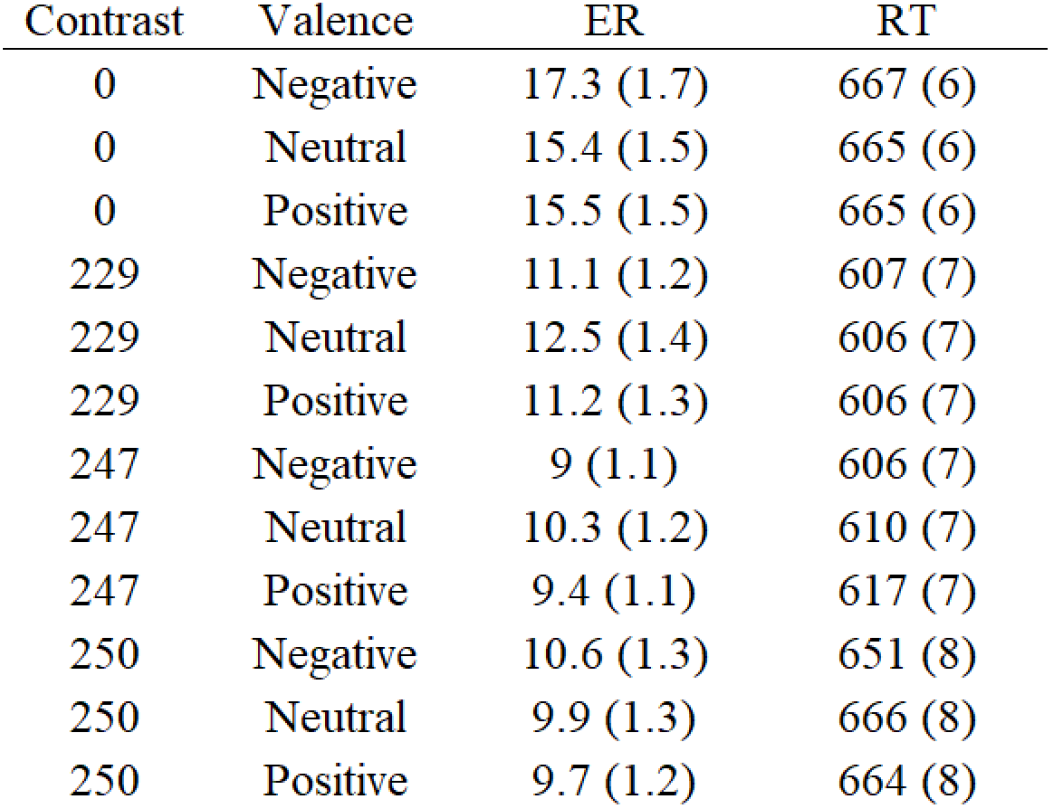
Means of ERs and RTs for each condition. Error rate (ER) and reaction time (RT) average values are reported as estimated marginal means from full GLMMs. ERs values were calculated (x 100) from estimated probabilities in the accuracy GLMMs. ( ) = standard error.

With respect to accuracy (shown as error rates to ease interpretation), the overall error rate (ER) was 11.8%. Least-square means for each condition are also shown in **Table 1** represented as ERs. The best GLMM included *Contrast* as a fixed effect, but not *Valence*. The *Contrast* main effect was significant [Wald χ2(3)=82.7, p<.001]. Corrected post-hoc z-tests revealed that ERs were highest in the 0 contrast (M=15,9%, SE=1.4%) [0 vs 229, z=5.5, p<.001; 0 vs 247, z=7.5, p<.001; 0 vs 250, z=7, p<.001] in comparison to the rest of contrasts (229, M=11,6%, SE=1.1%; 247, M=9,6%, SE=1%; 250, M=10,1%, SE=1.1%) [229 vs 247, z=2.2, p=.07; 229 vs 250, z=1.7, p=.16; 247 vs 250, z=.8, p=.413].

### 3.2. ERPs

The FMUT analysis over the ERP amplitudes for the 50-150 ms latency window yielded a significant *Valence* cluster [88-119 ms, 11 electrodes, Summed F = 230, cluster p = .039] corresponding to P1 (**Figure 2A**). A significant P1 cluster for the *Contrast x Valence* interaction was also found [88-127 ms, 26 electrodes, Summed F = 273, cluster p = .016] (**Figure 2D**). This cluster partially overlapped with the *Valence* cluster, but did not include all its electrodes, hence, we analyzed both clusters separately. Post-hoc Bonferroni corrected t-tests within the temporal window and electrodes of the *Valence* cluster indicated a significant *Negative > Positive* difference cluster [88-119 ms, 11 electrodes, Summed t= 171.4, cluster p <.001] (**Figure 2B** and **2C**). The post-hoc corrected t-tests for the *Contrast x Valence* interaction cluster indicated a significant *Negative > Positive* difference cluster only for the *250* contrast [88-127 ms, 23 electrodes, Summed t=400, cluster p <.001] (**Figure 2E** and **2F**). A *Contrast* cluster was also found [88-127 ms, 63 electrodes, Summed F = 7538, cluster p < .001].

**Figure 2.**
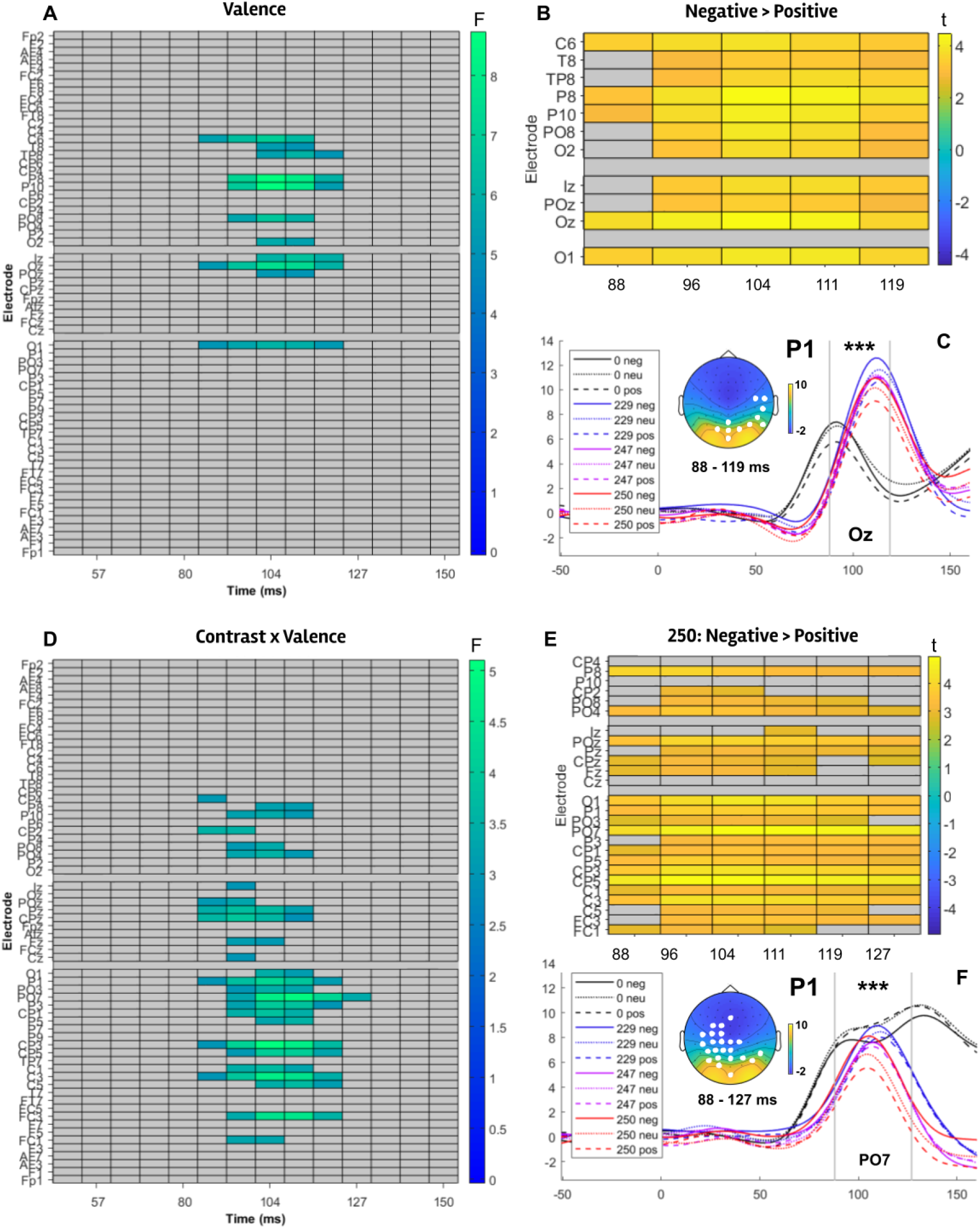
P1 negativity bias. **(A)** The Valence cluster for the 50-150 ms FMUT analysis is shown (blue-greenish rectangles represent the time window and electrodes). This cluster spanned 11 electrodes from 88 to 119 ms. **(B)** A significant post-hoc cluster revealed larger Negative than Positive P1 values in these time windows and electrodes (represented here as yellowish rectangles). **(C)** P1 amplitudes for each condition are shown for Oz (representative electrode). The average amplitudes and topographies for the cluster windows are shown. Significant electrodes are represented as white dots. **(D)** A significant Contrast x Valence cluster was found. This cluster spanned 26 electrodes from 88 to 127 ms. **(E)** A significant post-hoc cluster revealed larger Negative than Positive P1 values for the lowest contrast level (250). **(F)** P1 amplitudes for each condition are shown for PO7. Cluster p value <.001 = ^***^ .

Regarding the FMUT analysis on the ERP amplitudes within the P2-N2 window (150-600 ms), a significant *Valence* cluster was found (Figure 3), which corresponded to N2 and was widely distributed [213-525 ms, 56 electrodes, Summed F = 4498; cluster p =.005]. Post-hoc Bonferroni corrected t-tests within the temporal window and electrodes of the *Valence* cluster indicated two significant difference clusters *Positive > Negative* [213-354 ms, 39 electrodes, Summed t= 812, cluster p =.016] (Figure 4A and 4B) and Positive > Neutral [213-525 ms, 56 electrodes, Summed t= 2659, cluster p <.001] (Figure 4C and 4D). A Contrast cluster was also found [150-600 ms, 64 electrodes, Summed F = 114984, cluster p < .001].

**Figure 3.**
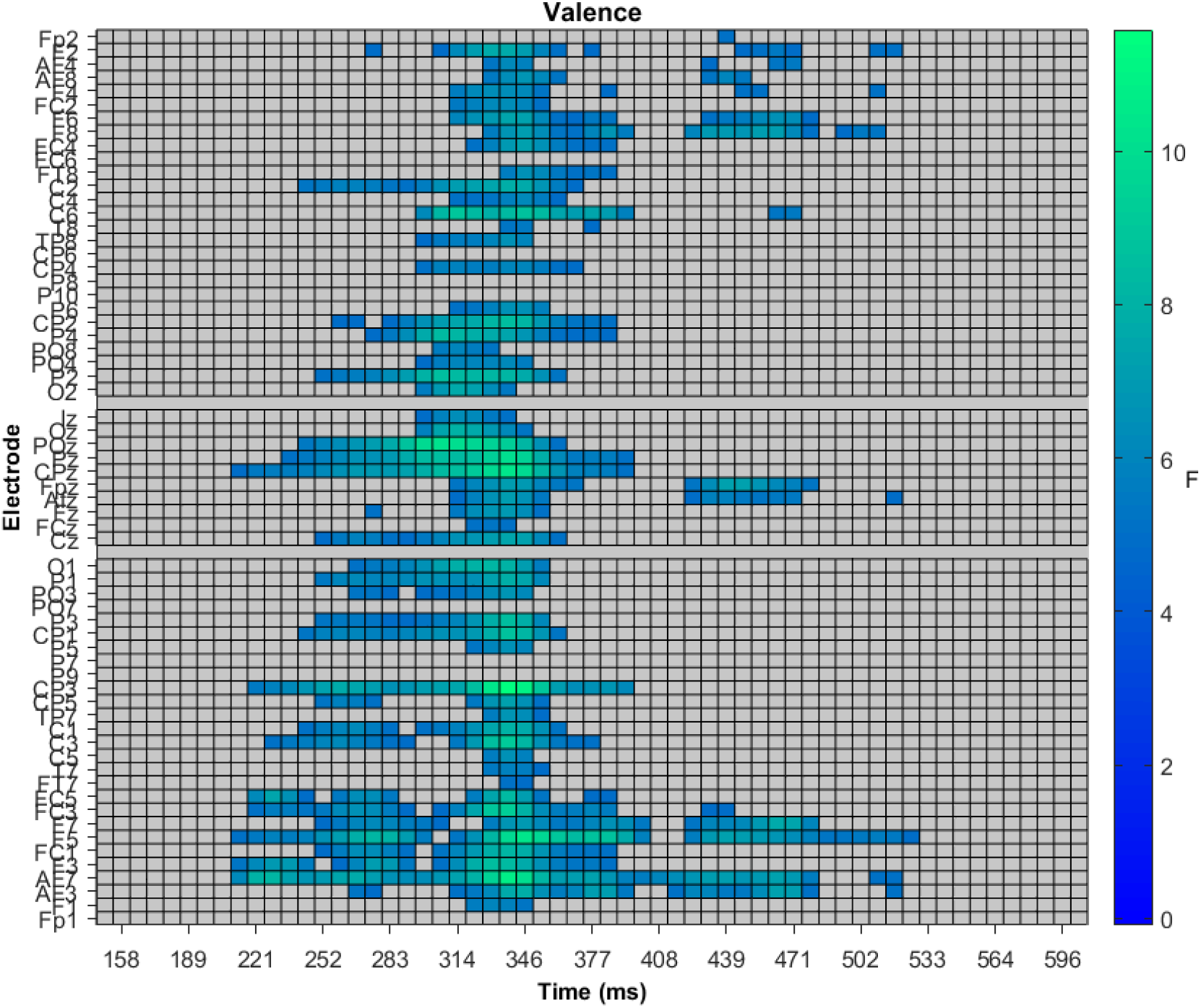
N2 valence cluster. The Valence cluster for the 150-600 ms FMUT analysis is shown. This cluster included 56 electrodes and ranged from 213 to 525 ms. Post-hoc contrasts within this Valence cluster are shown in Figure 4.

**Figure 4.**
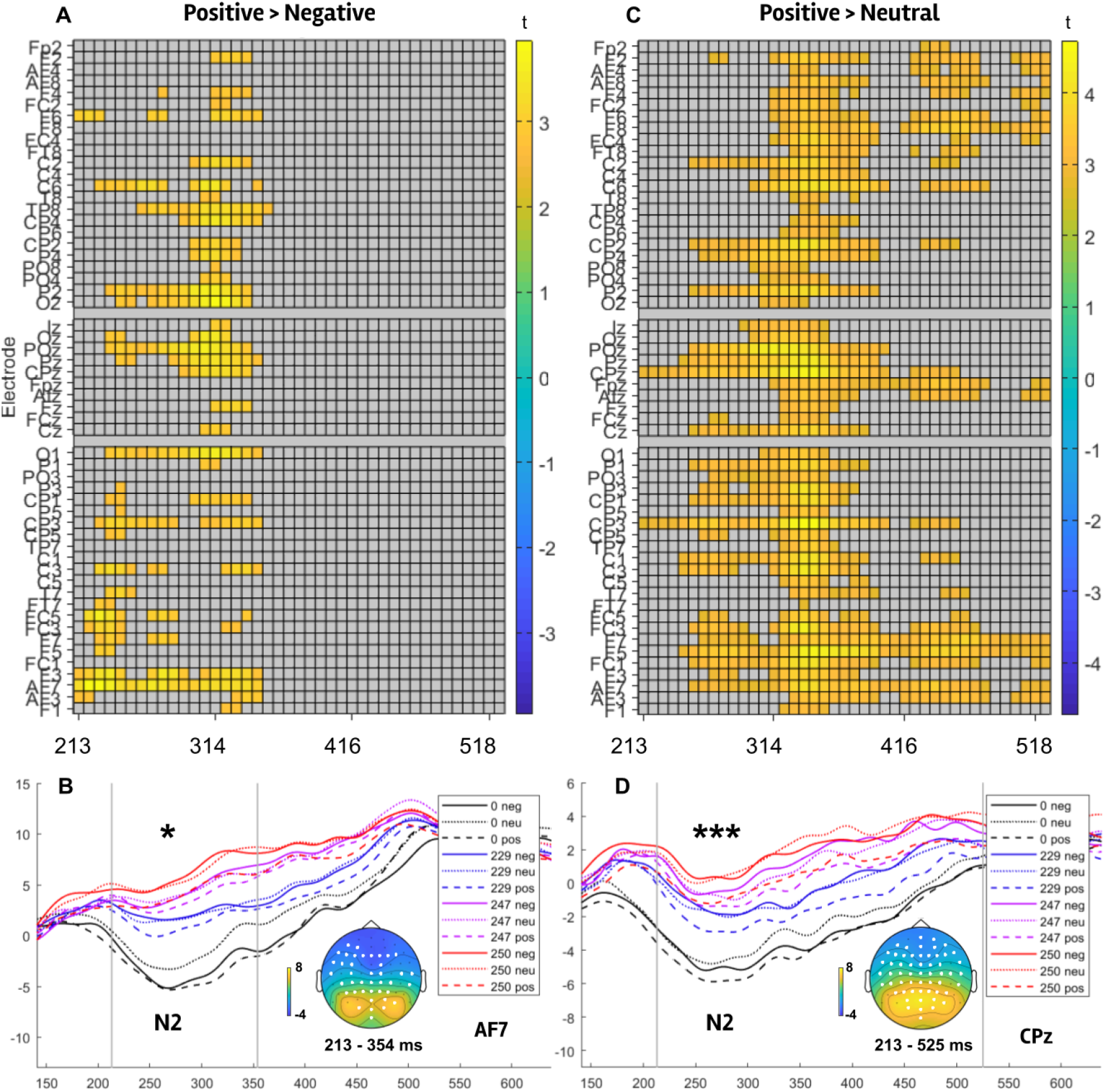
N2 positivity offset. The post-hoc contrasts within the Valence cluster (Figure 3) for the 150-600 ms FMUT analysis are shown. **(A)** A significant post-hoc cluster (39 electrodes; 213-354 ms) revealed larger (more negative) N2 values for Positive than Negative distractors. **(B)** N2 amplitudes for each condition are shown for AF7 (representative electrode). The average amplitudes and topographies for the cluster windows are shown. Significant electrodes are represented as white dots **(C)** A significant post-hoc cluster (56 electrodes; 213-525 ms) revealed larger N2 values for Positive than Neutral distractors. **(D)** N2 amplitudes for each condition are shown for CPz. Cluster p value: <.05 = ^*^ ; <.001 = ^***^ .

## 4. DISCUSSION

The objective of the present study was to investigate the influence of contrast level on exogenous attention to emotional scenes presenting different valences. To this aim, negative, neutral, and positive scenes were presented as distractors during a perceptual main task at four different contrast levels, while behavioral measures and ERPs were recorded. In short, at the behavioral level, we found that distractors were able to capture attention mainly when their contrast level was unaltered; at the neural level, negative distractors elicited larger P1 amplitudes, particularly for the lowest contrast level; additionally, regardless of the contrast level, positive scenes induced larger N2 amplitudes. These results will be discussed in more detail below.

At the behavioral level, reaction times and error rates indicate that, in general, distractors captured attention when their contrast level remained unaltered, and their emotional content did not significantly modulate the attentional capture as reflected in these variables. This is, the contrast degradation of distractors led to reduced errors and accelerated reaction times compared to unaltered scenes. An exception for this overall pattern was that, for the lowest contrast, reaction times were as slow as for intact distractors. This U-shaped pattern for reaction times might indicate a deeper assessment of the distractor in the lowest contrast condition, given its difficulty in being consciously perceived (less than 25% accurate identifications). Regarding the absence of effects by emotional valence of distractors on exogenous attention, even though meta-analytic data point to a bias towards negative and/ or positive stimuli (for a review see Carretié 2014), some previous studies have not found such an effect (e.g. De Cesarei et al., 2009; Holmes et al., 2003; Nordtröm & Wiens, 2012). Variability in crucial factors such as task nature (e.g., perceptual or digit categorization tasks), stimuli locations (e.g., at the center or at the periphery), and nature of the stimuli (e.g., faces or scenes) among others may underlie the absence of effects in some circumstances, as behavioral responses are single outputs which arise from a set of multiple cognitive stages such as perception, attention, decision making, and motor readiness, which may point in different directions (e.g., some inhibiting others). These discrete stages may be better disentangled by ERPs.

Regarding neural activity, and in chronological order, the initial ERP effect was observed in P1. First, a parieto-occipital P1 cluster (88-119 ms, 11 electrodes) revealed that, independently of the contrast level, negative emotional stimuli captured exogenous attention to a greater extent; specifically, amplitudes were larger for negative than for positive distractors. Second, a partially overlapping P1 cluster —albeit more spatially extensive—, reflecting an interaction between contrast and emotion, demonstrated that negative distractors elicited larger amplitudes than positive ones at the lowest contrast level (250 contrast: 88-127 ms, 23 electrodes). These findings evidence larger P1 amplitudes for negative than for positive stimuli. Specifically, the additional attentional effect only for the 250 contrast is likely explained by a partial habituation effect in P1 for the other contrast levels resulting from repeated exposure to emotional distractors across blocks. These P1 results are consistent with a negativity bias, which is the propensity to react or respond more strongly to negative than to positive stimuli (e.g., Cacioppo & Gardner, 1999; Cacioppo et al. 1999; Norris, 2021; Norris et al., 2010; Rozin & Royzman, 2001). In this context, this stronger early P1 response to negative distractors may evidence an evolutionary advantage, given that it is more critical for survival to fastly and accurately react to a life-threatening stimulus, than to approach or engage with positive stimuli. Relatedly, previous studies have found a negativity bias in P1 associated to an attentional capture of scenes (e.g., Carretié et al., 2004, 2016) or to the visual processing of images in general (emotional faces: Pourtois et al., 2004, 2005; emotional scenes: Smith et al., 2003, 2006). Crucially, the present findings further indicate that this negativity bias reflected in P1 operates even with poorly perceived negative distractors. Additionally, present results reinforce previous evidence showing P1 sensitivity to visual -non-emotional-stimuli at very low contrast levels (e.g. Ellemberg et al., 2001). These outcomes are consistent with those that point to a significant involvement of the magnocellular visual system in endogenous attention to emotional stimuli (e.g., Alorda et al., 2007; Miskovic et al., 2015; Pourtois et al., 2005; Vuilleumier et al., 2003), and, critically, with those that indicate a predominant role of the magnocellular pathway in attentional capture by negative stimuli (e.g., Carretié et al., 2012; 2017). In sum, our findings confirm a negativity bias reflected in P1, and they further reveal that it operates even at very low contrast levels (i.e., in conditions with poor visual perception). This suggests an evolutionary adaptation for fast magnocellular processing that signals hazardous settings.

Later on, N2 amplitudes also revealed an emotional effect, but of a different nature. Independently of the contrast level, positive distractor scenes elicited larger N2 amplitudes than negative (213-354 ms, 39 electrodes) and neutral ones (213-525 ms, 56 electrodes). Enhanced N2x amplitudes to positive distractors have already been reported in previous studies (scenes: e.g., Andreu et al., 2019; Carboni et al., 2017; Carretié et al., 2004; De Cesarei et al., 2009; but see Feng et al., 2012; on ADHD patients but not on controls, López-Martín et al., 2013; or to faces, Álvarez et al. 2022). However, the novel and crucial finding is that it is also elicited by scenes at very low contrast levels. Although the topography of the present N2 clusters is widely distributed throughout the scalp, the positive vs. neutral cluster extends for a long period at a frontal topography, consistent with previous findings indicating an anterior distribution for stronger positivity biases (e.g. Álvarez et al., 2022; Carboni et al., 2017; Carretié et al., 2004; López-Martín et al., 2013). In congruence with P1 amplitude results, N2 findings show that emotional visual stimuli are efficient in capturing attention in drastically impaired conditions and provide evidence of the role of magnocellular visual processing in exogenous attention. However, neural processes underlying P1 and N2 may be, at least in part, different. Assuming that emotional evaluation is not carried out by a single system nor at a single stage (e.g., Dixon et al., 2017), N2 reflects a bias for positive stimuli which might be interpreted in the context of a positivity offset. This refers to a bias towards pleasant stimuli (e.g., Cacioppo & Gardner, 1999; Cacioppo et al. 1999; Norris et al., 2010). This bias also has a relevant value for survival, given that it enables a person to approach novel stimuli, which encourages an exploratory behavior. In conjunction, P1 and N2 findings support the idea that it is crucial to detect first negative stimuli and then the positive ones (e.g., Carretié et al., 2004). Importantly, they further indicate that this detection occurs even in very low contrast conditions, lower than in previous studies. On a temporal scale, this suggests an adaptive magnocellular processing which prioritizes dangerous environments over satisfying ones.

In summary, the present study confirms that distractor scenes of negative valence capture attention at very early stages of processing (i.e., P1), and, as a novel and crucial finding, within very low contrast conditions, which indicates a predominant role of the magnocellular pathway. Positive distractors elicited an enhanced response at later latencies (i.e., N2) but, importantly, also independently of the contrast level. These results hallmark the efficiency of the brain in detecting first negative stimuli of high biological relevance and second stimuli with a positive valence. This temporal pattern evidences a magnocellular advantage for reacting to visual stimuli that may mean a significant risk to our survival, even without conscious awareness of their potential threat. Future research may extend the current findings by employing an inter-subject design. Intra-subject designs are the most recommended option for ERP studies in order to enhance statistical power while minimizing inter-subject variability (Luck, 2014). However, it is plausible that the intra-subject design combined with the blocked design, as employed here, led to a partial habituation of the P1 response to the remaining contrasts. As mentioned earlier, the blocked design was chosen because it facilitated the assessment of the lowest contrast levels without introducing contaminating influences from more visible images, which would have been present if contrast levels were randomized. Finally, gaining insight into whether fearful stimuli, as opposed to disgusting or anger-inducing stimuli (e.g., Krusemark & Li, 2011), or erotic stimuli compared to other appetizing stimuli (e.g., Feng et al., 2012), are responsible for shaping this pattern of biases would provide a deeper understanding of the role of these scenarios in guiding adaptive behavior under impaired visual conditions.

## SUPPLEMENTARY MATERIAL

**Table 1.**
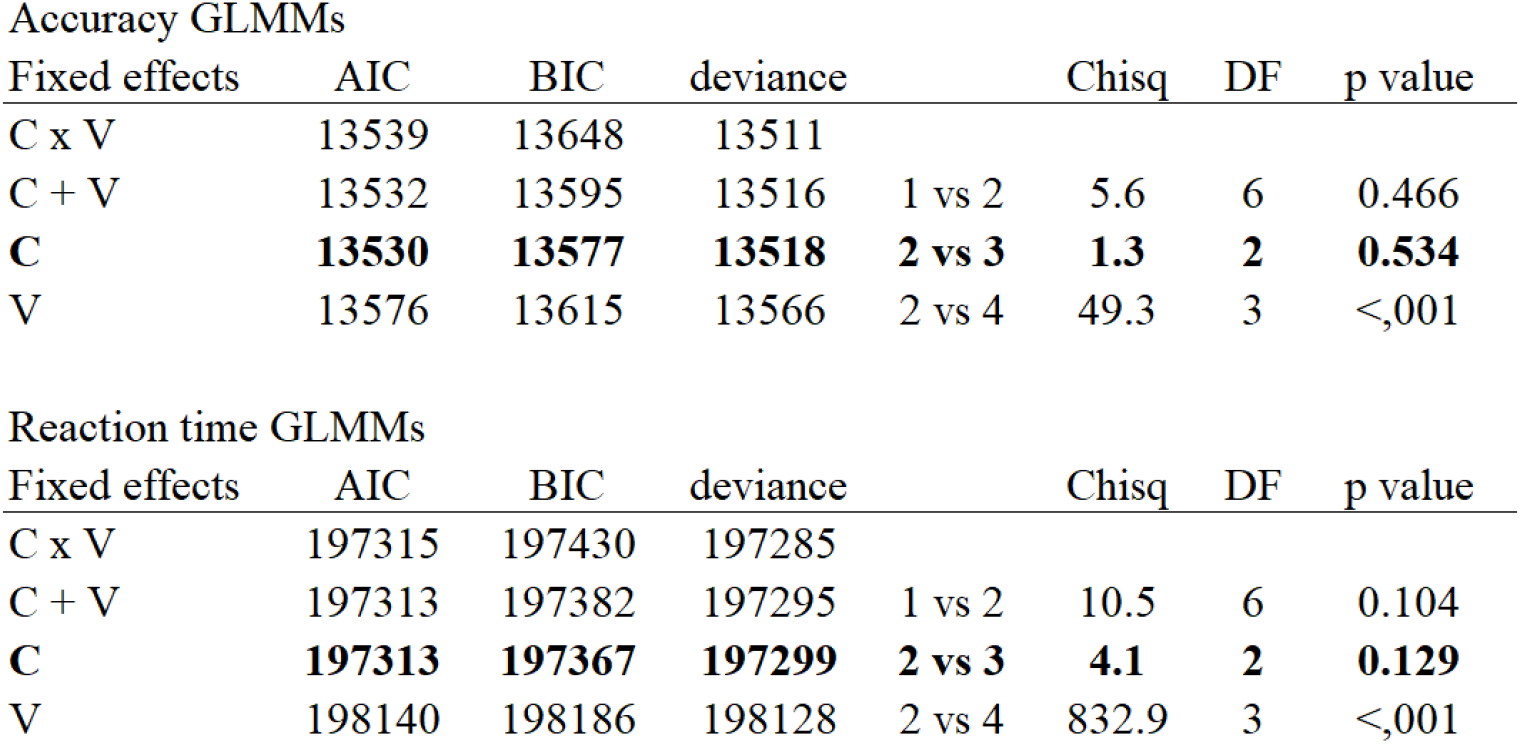
Generalized linear mixed model selection. GLMM model comparisons for different fixed effect structures are shown for accuracy and reaction time data. AIC= Akaike Information Criterion, BIC= Bayesian Information Criterion, Chisq = chi-square, DF= degrees of freedom.

Accuracy GLMMs
| Fixed effects | AIC | BIC | deviance |  | Chisq | DF | p value |
| --- | --- | --- | --- | --- | --- | --- | --- |
| C x V | 13539 | 13648 | 13511 |  |  |  |  |
| C + V | 13532 | 13595 | 13516 | 1 vs 2 | 5.6 | 6 | 0.466 |
| **C** | **13530** | **13577** | **13518** | **2 vs 3** | **1.3** | **2** | **0.534** |
| V | 13576 | 13615 | 13566 | 2 vs 4 | 49.3 | 3 | <.001 |

| Fixed effects | AIC | BIC | deviance |  | Chisq | DF | p value |
| --- | --- | --- | --- | --- | --- | --- | --- |
| C x V | 197315 | 197430 | 197285 |  |  |  |  |
| C + V | 197313 | 197382 | 197295 | 1 vs 2 | 10.5 | 6 | 0.104 |
| **C** | **197313** | **197367** | **197299** | **2 vs 3** | **4.1** | **2** | **0.129** |
| V | 198140 | 198186 | 198128 | 2 vs 4 | 832.9 | 3 | <.001 |

**Figure 1.**
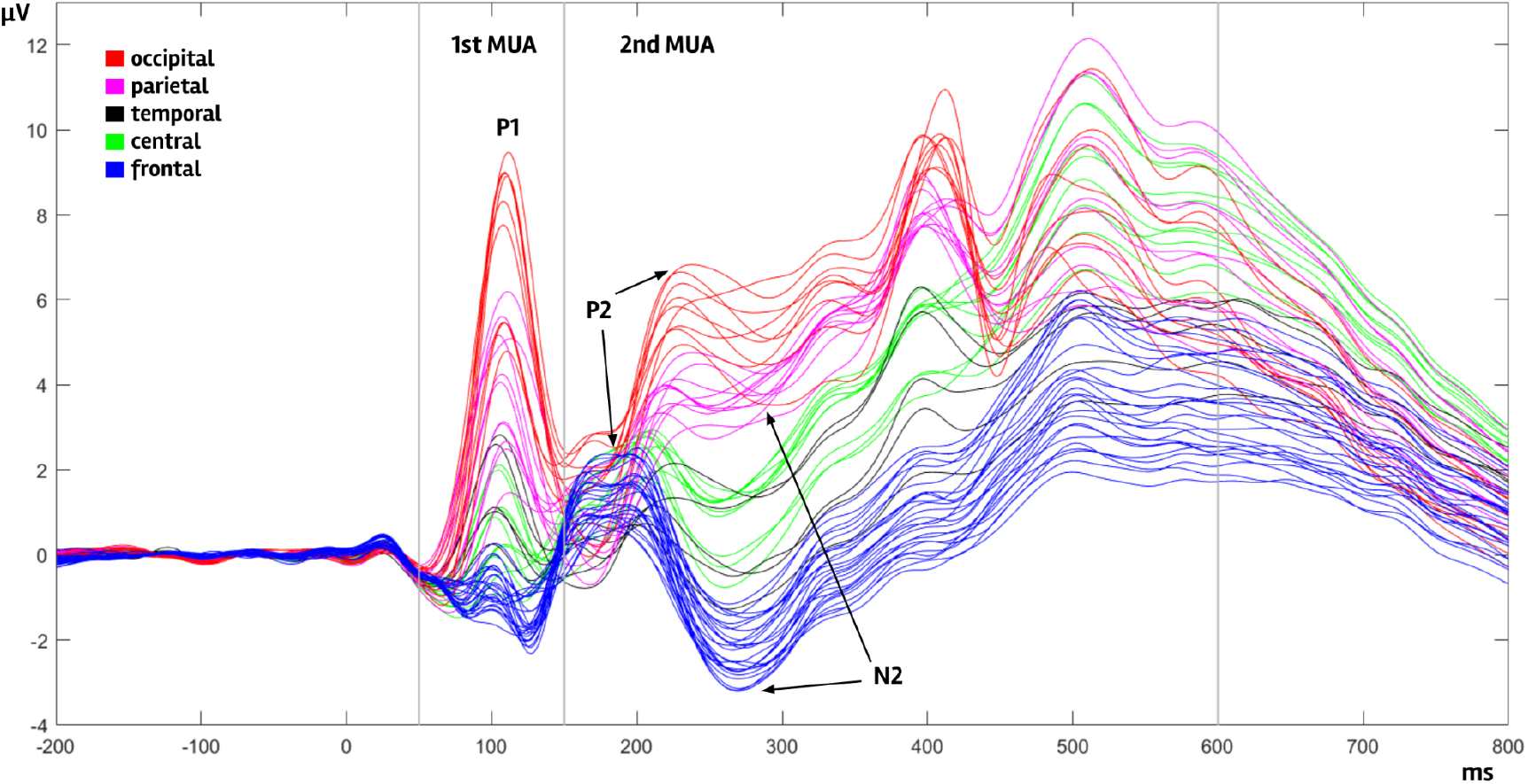
ERP average per channel. ERP average (across experimental conditions) for each of the 64 channels are shown, scalp regions are represented in different colors: red (occipital), magenta (parietal), black (temporal), green (central), and blue (frontal). Arrows point at P2 and N2 peaks at frontal and occipital regions. The two temporal windows selected for the Mass Univariate Analyses are shown (first: 50-150 ms, second: 150-600 ms). The occipito-parietal components within the 400-450 ms window correspond to the offset potential of the image.

## REFERENCES

Alorda, C., Serrano-Pedraza, I., Campos-Bueno, J. J., Sierra-Vázquez, V., & Montoya, P. (2007). Low spatial frequency filtering modulates early brain processing of affective complex pictures. Neuropsychologia, 45(14), 3223–3233.

Álvarez, F., Fernández-Folgueiras, U., Méndez-Bértolo, C., Kessel, D., & Carretié, L. (2022). Menstrual cycle and exogenous attention toward emotional expressions. Hormones and Behavior, 146, 105259.

Bates, D., Maechler, M., Bolker, B., Walker, S., Christensen, R. H. B., Singmann, H., … & Green, P. (2009). Package ‘lme4’. URL http://lme4.r-forge.r-project.org.

Brainard, D. H. (1997). The psychophysics toolbox. Spatial vision, 10(4), 433–436.

Buodo, G., Sarlo, M., & Munafo, M. (2010). The neural correlates of attentional bias in blood phobia as revealed by the N2pc. Social Cognitive and Affective Neuroscience, 5(1), 29–38.

Cacioppo, J. T., & Gardner, W. L. (1999). Emotion. Annual review of psychology, 50(1), 191–214.

Cacioppo, J. T., Gardner, W. L., & Berntson, G. G. (1999). The affect system has parallel and integrative processing components: Form follows function. Journal of personality and Social Psychology, 76(5), 839.

Carboni, A., Kessel, D., Capilla, A., & Carretié, L. (2017). The influence of affective state on exogenous attention to emotional distractors: behavioral and electrophysiological correlates. Scientific reports, 7(1), 1–13.

Carretié, L. (2014). Exogenous (automatic) attention to emotional stimuli: a review. Cognitive, Affective, & Behavioral Neuroscience, 14(4), 1228–1258.

Carretié, L., & Ruiz-Padial, E. (2016). Ambient light modulation of exogenous attention to threat. Brain topography, 29(6), 847–855.

Carretié, L., Albert, J., López-Martín, S., Hoyos, S., Kessel, D., Tapia, M., & Capilla, A. (2013a). Differential neural mechanisms underlying exogenous attention to peripheral and central distracters. Neuropsychologia, 51(10), 1838–1847.

Carretié, L., Fernández-Folgueiras, U., Álvarez, F., Cipriani, G. A., Tapia, M., & Kessel, D. (2022). Fast unconscious processing of emotional stimuli in early stages of the visual cortex. Cerebral Cortex, 32(19), 4331–4344.

Carretié, L., Hinojosa, J. A., Albert, J., López‐Martín, S., De La Gándara, B. S., Igoa, J. M., & Sotillo, M. (2008). Modulation of ongoing cognitive processes by emotionally intense words. Psychophysiology, 45(2), 188–196.

Carretié, L., Hinojosa, J. A., López-Martín, S., Albert, J., Tapia, M., & Pozo, M. A. (2009). Danger is worse when it moves: Neural and behavioral indices of enhanced attentional capture by dynamic threatening stimuli. Neuropsychologia, 47(2), 364–369.

Carretié, L., Hinojosa, J. A., López-Martín, S., & Tapia, M. (2007). An electrophysiological study on the interaction between emotional content and spatial frequency of visual stimuli. Neuropsychologia, 45(6), 1187–1195.

Carretié, L., Hinojosa, J. A., Martín‐Loeches, M., Mercado, F., & Tapia, M. (2004). Automatic attention to emotional stimuli: neural correlates. Human brain mapping, 22(4), 290–299.

Carretié, L., Hinojosa, J. A., Mercado, F., & Tapia, M. (2005). Cortical response to subjectively unconscious danger. Neuroimage, 24(3), 615–623.

Carretié, L., Kessel, D., Carboni, A., López-Martín, S., Albert, J., Tapia, M., … & Hinojosa, J. A. (2013b). Exogenous attention to facial vs non-facial emotional visual stimuli. Social Cognitive and Affective Neuroscience, 8(7), 764–773.

Carretié, L., Kessel, D., García-Rubio, M. J., Giménez-Fernández, T., Hoyos, S., & Hernández-Lorca, M. (2017). Magnocellular bias in exogenous attention to biologically salient stimuli as revealed by manipulating their luminosity and color. Journal of Cognitive Neuroscience, 29(10), 1699–1711.

Carretié, L., Ríos, M., Periáñez, J. A., Kessel, D., & Alvarez-Linera, J. (2012). The role of low and high spatial frequencies in exogenous attention to biologically salient stimuli. PloS one, 7(5), e37082.

Carretié, L., Ruiz-Padial, E., López-Martín, S., & Albert, J. (2011). Decomposing unpleasantness: Differential exogenous attention to disgusting and fearful stimuli. Biological psychology, 86(3), 247–253.

Carretié, L., Tapia, M., López-Martín, S., & Albert, J. (2019). EmoMadrid: An emotional pictures database for affect research. Motivation and Emotion, 43(6), 929–939.

Corbetta, M., Patel, G., & Shulman, G. L. (2008). The reorienting system of the human brain: From environment to theory of mind. Neuron, 58, 306–324.

De Cesarei, A., Codispoti, M., & Schupp, H. T. (2009). Peripheral vision and preferential emotion processing. Neuroreport, 20(16), 1439–1443.

Dixon, P. (2008). Models of accuracy in repeated-measures designs. Journal of Memory and Language, 59(4), 447–456.

Dixon, M. L., Thiruchselvam, R., Todd, R., & Christoff, K. (2017). Emotion and the prefrontal cortex: An integrative review. Psychological bulletin, 143(10), 1033.

Eimer, M., & Kiss, M. (2007). Attentional capture by task-irrelevant fearful faces is revealed by the N2pc component. Biological psychology, 74(1), 108–112.

Ekman, P. (1992). An argument for basic emotions. Cognition & emotion, 6(3-4), 169–200.

Ellemberg, D., Hammarrenger, B., Lepore, F., Roy, M. S., & Guillemot, J. P. (2001). Contrast dependency of VEPs as a function of spatial frequency: the parvocellular and magnocellular contributions to human VEPs. Spatial vision, 15(1), 99–111.

Feng, C., Wang, L., Wang, N., Gu, R., & Luo, Y. J. (2012). The time course of implicit processing of erotic pictures: An event-related potential study. Brain Research, 1489, 48–55.

Fernández-Folgueiras, U., Hernández-Lorca, M., Méndez-Bértolo, C., Álvarez, F., Giménez-Fernández, T., & Carretié, L. (2022). Exogenous Attention to Emotional Stimuli Presenting Realistic (3D) Looming Motion. Brain Topography, 35(5-6), 599–612.

Fields, E. C. (2017). Factorial mass univariate ERP toolbox. Computer software] Retrieved from https://github.com/ericcfields/FMUT/releases.

Fields, E. C., & Kuperberg, G. R. (2020). Having your cake and eating it too: Flexibility and power with mass univariate statistics for ERP data. Psychophysiology, 57(2), e13468.

Groppe, D. M., Urbach, T. P., & Kutas, M. (2011). Mass univariate analysis of event‐related brain potentials/fields I: A critical tutorial review. Psychophysiology, 48(12), 1711–1725.

Hinojosa, J. A., Carretié, L., Valcárcel, M. A., Méndez-Bértolo, C., & Pozo, M. A. (2009). Electrophysiological differences in the processing of affective information in words and pictures. Cognitive, Affective, & Behavioral Neuroscience, 9(2), 173–189.

Holmes, A., Vuilleumier, P., & Eimer, M. (2003). The processing of emotional facial expression is gated by spatial attention: evidence from event-related brain potentials. Cognitive Brain Research, 16(2), 174–184.

Jung, T. P., Makeig, S., Westerfield, M., Townsend, J., Courchesne, E., & Sejnowski, T. J. (2000). Removal of eye activity artifacts from visual event-related potentials in normal and clinical subjects. Clinical Neurophysiology, 111(10), 1745–1758.

Kleiner, M., Brainard, D., & Pelli, D. (2007). What’s new in Psychtoolbox-3?.

Krusemark, E. A., & Li, W. (2011). Do all threats work the same way? Divergent effects of fear and disgust on sensory perception and attention. Journal of Neuroscience, 31(9), 3429–3434.

Lee, K. Y., Lee, T. H., Yoon, S. J., Cho, Y. S., Choi, J. S., & Kim, H. T. (2010). Neural correlates of top–down processing in emotion perception: an ERP study of emotional faces in white noise versus noise-alone stimuli. Brain Research, 1337, 56–63.

Lo, S., & Andrews, S. (2015). To transform or not to transform: Using generalized linear mixed models to analyse reaction time data. Frontiers in psychology, 6, 1171.

López-Martín, S., Albert, J., Fernández-Jaén, A., & Carretié, L. (2013). Emotional distraction in boys with ADHD: Neural and behavioral correlates. Brain and cognition, 83(1), 10–20.

Mavratzakis, A., Herbert, C., & Walla, P. (2016). Emotional facial expressions evoke faster orienting responses, but weaker emotional responses at neural and behavioural levels compared to scenes: A simultaneous EEG and facial EMG study. Neuroimage, 124, 931–946.

Mercado, F., Carretié, L., Hinojosa, J. A., & Penacoba, C. (2009). Two successive phases in the threat‐related attentional response of anxious subjects: neural correlates. Depression and anxiety, 26(12), 1141–1150.

Mercado, F., Carretié, L., Tapia, M., & Gómez-Jarabo, G. (2006). The influence of emotional context on attention in anxious subjects: neurophysiological correlates. Journal of anxiety disorders, 20(1), 72–84.

Merigan, W. H., & Maunsell, J. H. (1993). How parallel are the primate visual pathways?. Annual review of neuroscience.

Miskovic, V., Martinovic, J., Wieser, M. J., Petro, N. M., Bradley, M. M., & Keil, A. (2015). Electrocortical amplification for emotionally arousing natural scenes: The contribution of luminance and chromatic visual channels. Biological psychology, 106, 11–17.

Norris, C. J. (2021). The negativity bias, revisited: Evidence from neuroscience measures and an individual differences approach. Social neuroscience, 16(1), 68–82.

Nordström, H., & Wiens, S. (2012). Emotional event-related potentials are larger to figures than scenes but are similarly reduced by inattention. BMC neuroscience, 13, 1–10.

Norris, C. J., Gollan, J., Berntson, G. G., & Cacioppo, J. T. (2010). The current status of research on the structure of evaluative space. Biological psychology, 84(3), 422–436.

Öhman, A., Hamm, A., & Hugdahl, K. (2000). Cognition and the autonomic nervous system: Orienting, anticipation, and conditioning. In J. T. Cacioppo, L. G. Tassinary, & G. G. Bernston (Eds.), Handbook of psychophysiology (2nd ed., pp. 533–575). Cambridge: Cambridge University Press.

Oostenveld, R., Fries, P., Maris, E., & Schoffelen, J. M. (2011). FieldTrip: open source software for advanced analysis of MEG, EEG, and invasive electrophysiological data. Computational intelligence and neuroscience, 2011.

Pourtois, G., Dan, E. S., Grandjean, D., Sander, D., & Vuilleumier, P. (2005). Enhanced extrastriate visual response to bandpass spatial frequency filtered fearful faces: Time course and topographic evoked‐potentials mapping. Human brain mapping, 26(1), 65–79.

Pourtois, G., Grandjean, D., Sander, D., & Vuilleumier, P. (2004). Electrophysiological correlates of rapid spatial orienting towards fearful faces. Cerebral cortex, 14(6), 619–633.

Purpura, K., Kaplan, E., & Shapley, R. M. (1988). Background light and the contrast gain of primate P and M retinal ganglion cells. Proceedings of the National Academy of Sciences, 85(12), 4534–4537.

Rozin, P., & Royzman, E. B. (2001). Negativity bias, negativity dominance, and contagion. Personality and social psychology review, 5(4), 296–320.

Schönwald, L. I., & Müller, M. M. (2014). Slow biasing of processing resources in early visual cortex is preceded by emotional cue extraction in emotion–attention competition. Human Brain Mapping, 35(4), 1477–1490.

Soares, S. C., Kessel, D., Hernández-Lorca, M., García-Rubio, M. J., Rodrigues, P., Gomes, N., & Carretié, L. (2017). Exogenous attention to fear: Differential behavioral and neural responses to snakes and spiders. Neuropsychologia, 99, 139–147.

Smith, N. K., Cacioppo, J. T., Larsen, J. T., & Chartrand, T. L. (2003). May I have your attention, please: Electrocortical responses to positive and negative stimuli. Neuropsychologia, 41(2), 171–183.

Smith, N. K., Larsen, J. T., Chartrand, T. L., Cacioppo, J. T., Katafiasz, H. A., & Moran, K. E. (2006). Being bad isn’t always good: affective context moderates the attention bias toward negative information. Journal of personality and social psychology, 90(2), 210.

Sussman, T. J., Heller, W., Miller, G. A., & Mohanty, A. (2013). Emotional distractors can enhance attention. Psychological science, 24(11), 2322–2328.

Team, R. C. (2013). R: A language and environment for statistical computing.

Thom, N., Knight, J., Dishman, R., Sabatinelli, D., Johnson, D. C., & Clementz, B. (2014). Emotional scenes elicit more pronounced self-reported emotional experience and greater EPN and LPP modulation when compared to emotional faces. Cognitive, Affective, & Behavioral Neuroscience, 14, 849–860.

Tootell, R. B., & Nasr, S. (2017). Columnar segregation of magnocellular and parvocellular streams in human extrastriate cortex. Journal of Neuroscience, 37(33), 8014–8032.

Tootell, R. B., Hamilton, S. L., & Switkes, E. (1988). Functional anatomy of macaque striate cortex. IV. Contrast and magno-parvo streams. Journal of Neuroscience, 8(5), 1594–1609.

Trauer, S. M., Andersen, S. K., Kotz, S. A., & Müller, M. M. (2012). Capture of lexical but not visual resources by task-irrelevant emotional words: A combined ERP and steady-state visual evoked potential study. Neuroimage, 60(1), 130–138.

Vuilleumier, P., Armony, J. L., Driver, J., & Dolan, R. J. (2003). Distinct spatial frequency sensitivities for processing faces and emotional expressions. Nature neuroscience, 6(6), 624–631.

Whelan, R. (2008). Effective analysis of reaction time data. The psychological record, 58(3), 475–482.

Zuur, A. F., Ieno, E. N., Walker, N. J., Saveliev, A. A., & Smith, G. M. (2009). Mixed effects models and extensions in ecology with R. New York: springer.

